# Post-reactivation mifepristone impairs generalisation of strongly-conditioned contextual fear memories

**DOI:** 10.1101/2020.05.11.088922

**Authors:** Charlotte R. Flavell, Rebecca M. Gascoyne, Jonathan L. C. Lee

**Affiliations:** University of Birmingham, School of Psychology, Edgbaston, Birmingham B15 2TT; Aston University, Birmingham B4 7ET

**Keywords:** reconsolidation, context, fear, mifepristone, glucocorticoid

## Abstract

The efficacy of pharmacological disruption of fear memory reconsolidation depends on several factors, including memory strength and age. We built on previous observations that systemic treatment with the nootropic nefiracetam potentiates cued fear memory destabilization to facilitate mifepristone-induced reconsolidation impairment. Here, we applied nefiratecam and mifepristone to strongly-conditioned, 1-week old contextual fear memories in male rats. Unexpectedly, the combined treatment did not result in impairment of contextual fear expression. However, mifepristone did reduce freezing to a novel context. These observations suggest that strong and established contextual fear memories do undergo destabilization without the need for pharmacological facilitation, and that impairments in strong context fear memory reconsolidation can manifest as a reduction in generalization.

---

The disruption of fear memory reconsolidation may present an opportunity to diminish maladaptive memories in conditions such as PTSD (Paulus et al. 2019). Multiple pharmacological agents have been identified, which when administered around the time of memory reactivation (typically through cue re-exposure) have been shown to impair subsequent fear memory expression (Reichelt and Lee 2013; Bolsoni and Zuardi 2019). One such drug is the glucocorticoid receptor antagonist mifepristone (Jin et al. 2007; Pitman et al. 2011; Flavell and Lee 2019).

The efficacy of reconsolidation-blocking treatments to disrupt subsequent memory expression is dependent upon the success of the reactivation session in destabilizing the target memory (Dudai 2012). Memory destabilisation can be blocked pharmacologically (Ben Mamou et al. 2006; Wideman et al. 2018), and there are multiple boundary conditions on memory reconsolidation, which describe parametric conditions under which memory reactivation fails to destabilise the memory (Lee 2009; Haubrich and Nader 2018). One such boundary condition that is particularly relevant to PTSD is that of memory strength; stronger fear memories are generally more difficult to destabilize (Wang et al. 2009), requiring more extensive cue re-exposure (Suzuki et al. 2004).

In order to avoid extensive parametric experimentation, which would not suit clinical intervention, the ability to enhance memory destabilization pharmacologically is of potential benefit. The stimulation of fear memory destabilization has been observed under a number of different settings and using a variety of pharmacological treatments (Bustos et al. 2010; Lee and Flavell 2014; Gazarini et al. 2015; Flavell and Lee 2019). We have recently demonstrated that the nootropic nefiracetam can enhance the destabilization of cued fear memories, which facilitates the impairment of reconsolidation by mifepristone (Flavell and Lee 2019). This enhancement of destabilization was observed under conditions of relatively mild fear conditioning (a single footshock), albeit resulting in high levels of cued freezing. Therefore, it remains unclear whether the facilitative effect of nefiracetam translates to paradigms that perhaps more closely replicate the clinical condition of PTSD, in which the fearful/traumatic memory is substantially stronger.

Here, we applied the combined nefiracetam-mifepristone treatment to a strong contextual fear memory paradigm consisting of 10 footshocks. *A priori* we aimed to implement a variation of the stress-enhanced fear learning paradigm (Rau et al. 2005), but found that the initially-conditioned contextual fear generalised substantially to the second context. Nevertheless, we observed differences between the groups on that generalised context fear expression, indicating contrary to our predictions a direct effect of mifepristone in the absence of nefiracetam treatment.

40 male Lister Hooded rats (Charles River, UK; 200-225 g at the start of the experiment) were housed in quads under a 12 h light/dark cycle (lights on at 0700) at 21°C with food and water provided ad libitum apart from during the behavioural sessions. Standard cages contained aspen chip bedding and environmental enrichment was available in the form of a Plexiglass tunnel. Experiments took place in a behavioural laboratory between 0830 and 1300. At the end of the experiment, animals were humanely killed via a rising concentration of CO2; death was confirmed by cervical dislocation. Principles of laboratory animal care were followed, as approved by the University of Birmingham Animal Welfare and Ethical Review Body and in accordance to the United Kingdom Animals (Scientific Procedures) Act 1986, Amendment Regulations 2012 (PPL P3B19D9B2).

Rats were initially fear conditioned in Context A (CXA; MedAssociates [VT] chamber [ENV-008] with triangular insert [ENV-008-IRT], viewing window in sound-attenuating chamber [ENV-022MD-WF], floorbars with alternating diameters [VFC-005-L], and 3 drops of 10% acetic acid). Ten 0.7-mA, 1-s footshocks were delivered in a 60-min session with an inter-trial interval averaging 5 min (range 150 – 450 s). 7 days later, the rats were returned to CXA for a 5-min reactivation session. All drugs were administered systemically at previously-established doses and timepoints (Flavell and Lee 2019). Nefiracetam (Sigma, UK) was injected at 3 mg/kg (6 mg/ml in saline, i.p.) 1 hr before reactivation. Mifepristone (Generon, UK) was injected at 30 mg/kg (60 mg/ml in propylene glycol, s.c.) immediately after memory reactivation. Allocation to drug treatment was fully randomised within each experimental cohort of 8 rats. On the day after reactivation, the rats were again returned to CXA for a 5-min test session.

In CXA, there was little evidence for an effect of Nefiracetam or Mifepristone at reactivation or test. Freezing was scored manually at 5-s intervals by an experimenter blind to the experimental status of the animals. Repeated-measures ANOVA (JASP Team 2016) revealed that there was neither an overall effect of Nefiracetam (F(1,27)=1.25, p=0.27, η^2^_p_=0.044, BF_inc_=0.38), nor a Nefiracetam x Session interaction (F(1,27)=0.014, p=0.91, η^2^_p_=0.001, BF_inc_=0.35). Similarly, there was no overall effect of Mifepristone (F(1,27)=2.84, p=0.10, η^2^_p_=0.095, BF_inc_=0.74), nor a Mifepristone x Session interaction (F(1,27)=1.08, p=0.31, η^2^_p_=0.039, BF_inc_=0.68). Finally, there was no evidence for an interaction between Nefiracetam and Mifepristone (F(1,27)=0.042, p=0.84, η^2^_p_=0.002, BF_inc_=0.31; Nefiracetam x Mifepristone x Session: F(1,27)=0.65, p=0.43, η^2^_p_=0.023, BF_inc_=0.10). Planned analyses of simple main effects revealed little evidence for an effect of Mifepristone under any condition (Fig. 1A). Therefore, when tested in the conditioning context, there was no evidence for a disruptive effect of peri-reactivation Nefiracetam and/or Mifepristone. Across drug groups, freezing declined from the reactivation session to test (Main effect of Session: F(1,27)=35.7, p<0.001, η^2^_p_=0.57, BF_inc_=16557).

**Fig. 1.**
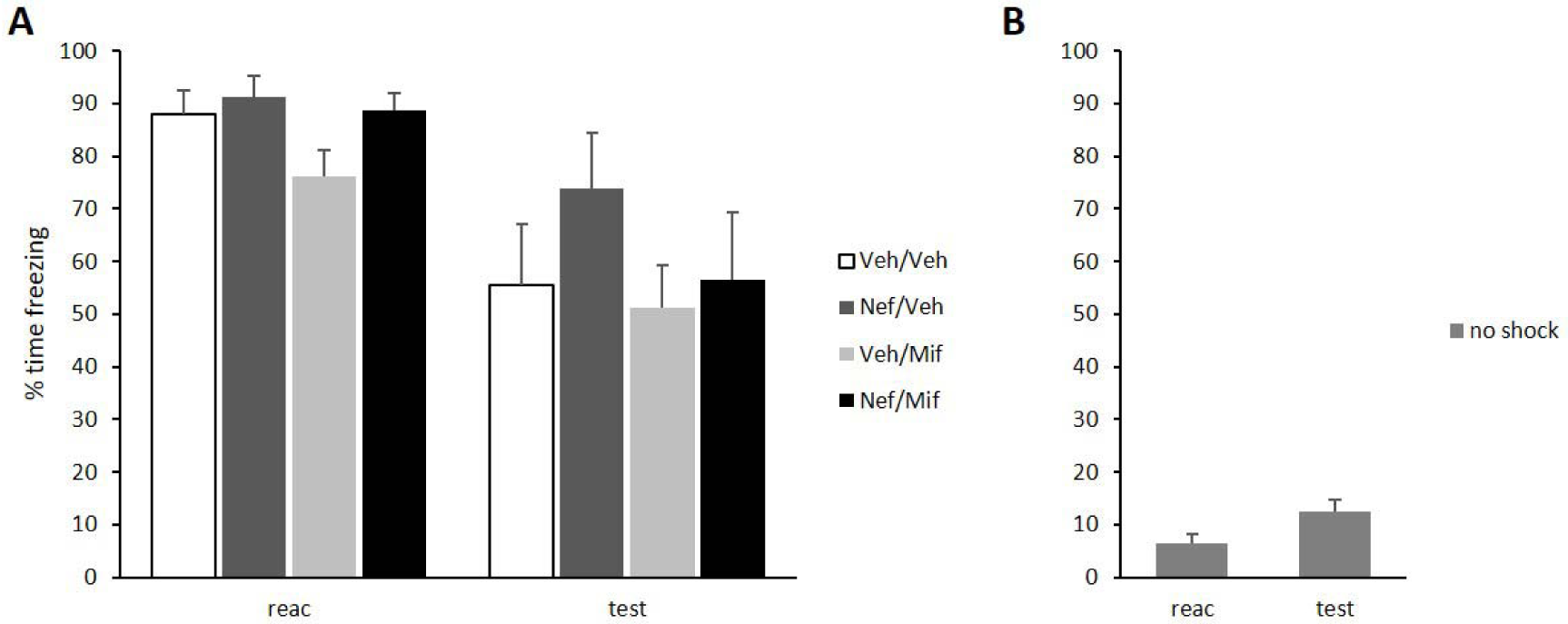
Conditioned freezing to Context A at reactivation (reac) and test in groups that received drug administration (A) and the non-conditioned no shock control (B). Peri-reactivation nefiracetam (Nef) and mifepristone (Mif) had no obvious effect on freezing in either session. Planned analyses of the effect of Mifepristone showed no drug effects (reactivation: vehicle p=0.094, nefiracetam p=0.610; test: vehicle p=0.340, nefiracetam p=0.275). Data presented as mean + SEM (n = 8 per group).

When comparing each group to a no shock non-conditioned control that was exposed to CXA instead of being fear conditioned, freezing was consistently higher at both reactivation and test (Fig. 1B). There was a Group x Session interaction (F(4,34)=4.09, p=0.008, η^2^_p_=0.33, BF_inc_=24.6), as well as a main effect of Group (F(4,34)=24.4, p<0.001, η^2^_p_=0.74, BF_inc_=4.8 × 10^7^). Analysis of simple main effects revealed Group differences at both reactivation at test (p’s<0.001). Post-hoc Tukey-corrected pairwise comparisons on the main effect of Group across both session showed that the non-conditioned group froze lower than each of the other groups (p’s<0.001, BF10’s>4.3×10^6^), which did not differ from each other (p’s>18s, BF10’s<0.78 [apart from 3.01 for Nef/Veh vs Veh/Mif]).

6 days after the CXA test, the rats were placed into a different Context B (CXB; MedAssociates [VT] chamber [ENV-008] with no insert or viewing window in the sound-attenuating chamber [ENV-018MD], standard equal floorbars [ENV-005], no added odor, and a video camera mounted visibly above the chamber), for a 3-min session, with delivery of a 0.4-mA, 2-s footshock after 120 s. In CXB, freezing behaviour during the conditioning session (quantified automatically by videotracking software; Viewpoint Lifesciences) revealed evidence for a disruptive effect of Mifepristone in the absence of Nefiracetam in the pre-shock period. The overall analysis revealed neither an overall effect of Nefiracetam (F(1,28)=1.60, p=0.22, η^2^_p_=0.054, BF_inc_=0.71), nor a Nefiracetam x Phase interaction (F(1,28)=3.0, p=0.10, η^2^_p_=0.096, BF_inc_=1.1). Similarly, there was no overall effect of Mifepristone (F(1,28)=1.94, p=0.17, η^2^_p_=0.065, BF_inc_=0.57), nor a Mifepristone x Phase interaction (F(1,28)=0.042, p=0.84, η^2^_p_=0.001, BF _inc_=0.48). Finally, there was no evidence for an interaction between Nefiracetam and Mifepristone (F(1,28)=1.63, p=0.21, η^2^_p_=0.055, BF_inc_=0.60; Nefiracetam x Mifepristone x Phase: F(1,28)=1.54, p=0.23, η^2^_p_=0.052, BF_inc_=0.29). However, planned analyses of simple main effects revealed evidence for an effect of Mifepristone only in rats previously injected with vehicle (not Nefiracetam) and only in the pre-shock period (Fig. 2A). Therefore, injection of Mifepristone without Nefiracetam pre-treatment reduced freezing to CXB prior to conditioning, but did not appear to affect freezing after footshock delivery in CXB.

**Fig. 2.**
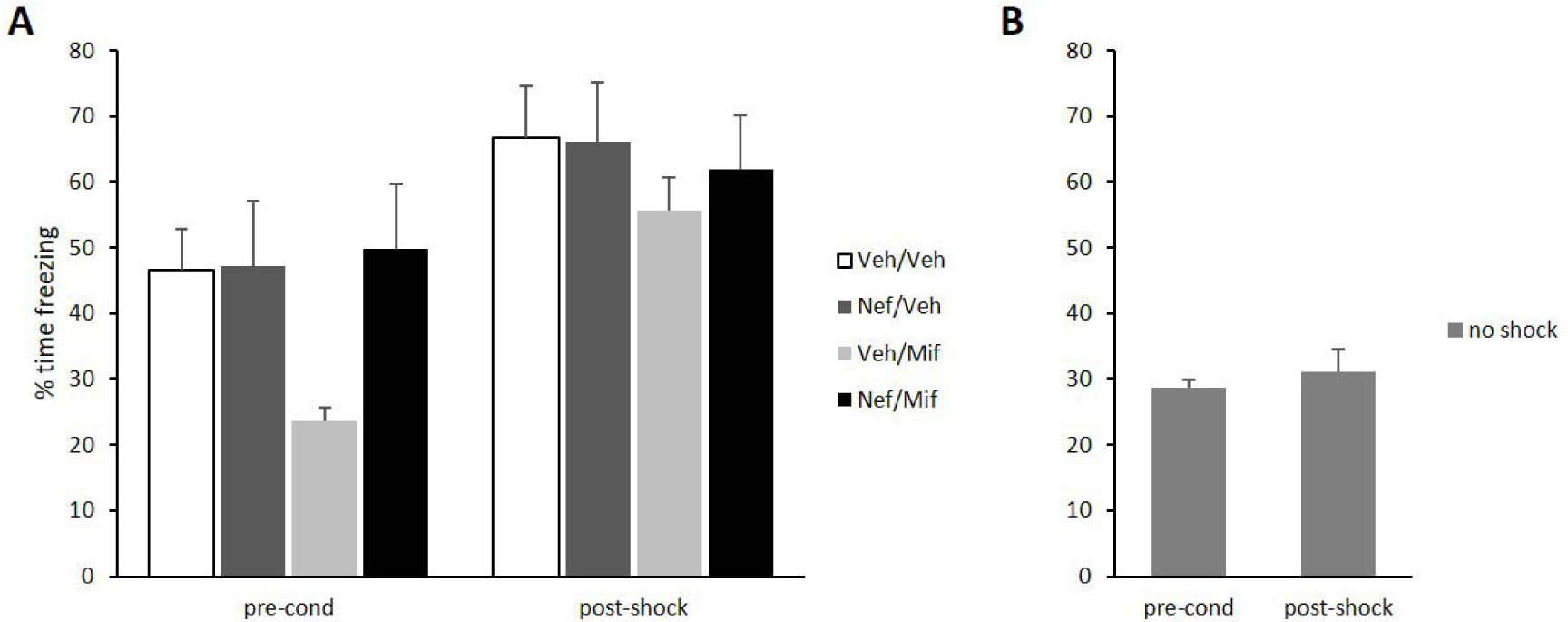
Conditioned freezing to Context B prior to (pre-cond) and after (post-shock) footshock delivery in groups that previously received drug administration (A) and the non-conditioned no shock control (B). Mifepristone (Mif) injected immediately after Context A memory reactivation resulted in impaired freezing to Context B in the pre-shock period. This was not observed when reactivation was preceded by nefiracetam (Nef) injection. Planned analyses of the effect of Mifepristone showed selective drug effects (pre-cond: vehicle p=0.003, nefiracetam p=0.837; post-shock: vehicle p=0.196, nefiracetam p=0.717). Data presented as mean + SEM (n = 8 per group).

When comparing each group to the non-conditioned control, freezing was consistently higher at both reactivation and test in all groups expect the Vehicle-Mifepristone group (Fig. 2B). There was a Group x Phase interaction (F(4,35)=3.30, p=0.022, η^2^_p_=0.27, BF_inc_=10.4), as well as a main effect of Group (F(4,35)=4.62, p=0.004, η^2^_p_=0.35, BF_inc_=30.2). Analysis of simple main effects revealed Group differences both prior to and after footshock delivery (p’s<0.021). Post-hoc pairwise comparisons on the main effect of Group across both session showed that the non-conditioned group froze lower than each of the other groups (p’s<0.025, BF_10_’s>69), apart from the Vehicle-Mifepristone group (p=0.97, BF_10_=1.39). However, the Vehicle-Mifepristone group did not differ from the other groups (p’s>0.29, BF_10_’s<2.63). Given the apparent impairment in the Vehicle-Mifepristone group in the pre-shock period, we conducted an exploratory ANCOVA to determine whether any reduction in freezing during the post-shock period might be due to any additional effect. This analysis showed a significant difference between the non-conditioned control and all other groups (p<0.035, BF_10_’s>21), including the Vehicle-Mifepristone group (p=0.010, BF_10_=101), but not the Nefiracetam-Mifepristone group (p=0.172, but BF_10_=21.2). Therefore, there was little evidence for a disruption of conditioning in CXB.

As the pre-shock period in CXB was 2 min, compared to the 5-min reactivation and test sessions in CXA, we re-analysed the first 2 min of the CXA sessions (Table 1). The statistical patterns were not different to those for the full 5 min of the sessions.

**Table 1.** Mean ± SEM freezing in the first 2 min of the reactivation and test sessions in Context A.

| Group | Reactivation | Test |
| --- | --- | --- |
| Vehicle/Vehicle | 79.2 ± 5.8 | 50.0 ± 8.9 |
| Nefiracetam/Vehicle | 86.5 ± 6.6 | 66.1 ± 9.6 |
| Vehicle/Mifepristone | 72.4 ± 7.0 | 40.6 ± 10.4 |
| Nefiracetam/Mifepristone | 84.9 ± 4.1 | 58.9 ± 13.4 |
| No shock | 4.7 ± 2.6 | 13.5 ± 3.9 |

On the day after conditioning in CXB, the rats were returned to CXB for a 30-min test session. At the CXB test, there was some evidence for an effect of Mifepristone to reduce freezing across the session, particularly with Vehicle pre-treatment (Fig. 3A). There was neither an overall effect of Nefiracetam (F(1,28)=0.49, p=0.49, η^2^_p_=0.017, BF_inc_=0.19), nor a Nefiracetam x Bin interaction (F(2.5,71.2)=0.74, p=0.51, η^2^_p_=0.026, BF_inc_=0.070). However, there was an overall effect of Mifepristone (F(1,28)=5.67, p=0.024, η^2^_p_=0.17, BF_inc_=1.08), but no Mifepristone x Bin interaction (F(2.5,71.2)=0.53, p=0.64, η^2^_p_=0.018, BF_inc_=0.12). Finally, there was no evidence for an interaction between Nefiracetam and Mifepristone (F(1,28)=1.09, p=0.31, η^2^_p_=0.038, BF_inc_=0.23; Nefiracetam x Mifepristone x Bin: F(2.5,71.2)=0.33, p=0.77, η^2^_p_=0.012, BF_inc_=0.002). Planned analyses of simple main effects revealed more evidence for an effect of Mifepristone in rats pre-treated with Vehicle (p=0.031) than those administered Nefiracetam (p<0.36). Therefore, the deficit observed prior to conditioning in CXB was not eliminated by conditioning.

**Fig. 3.**
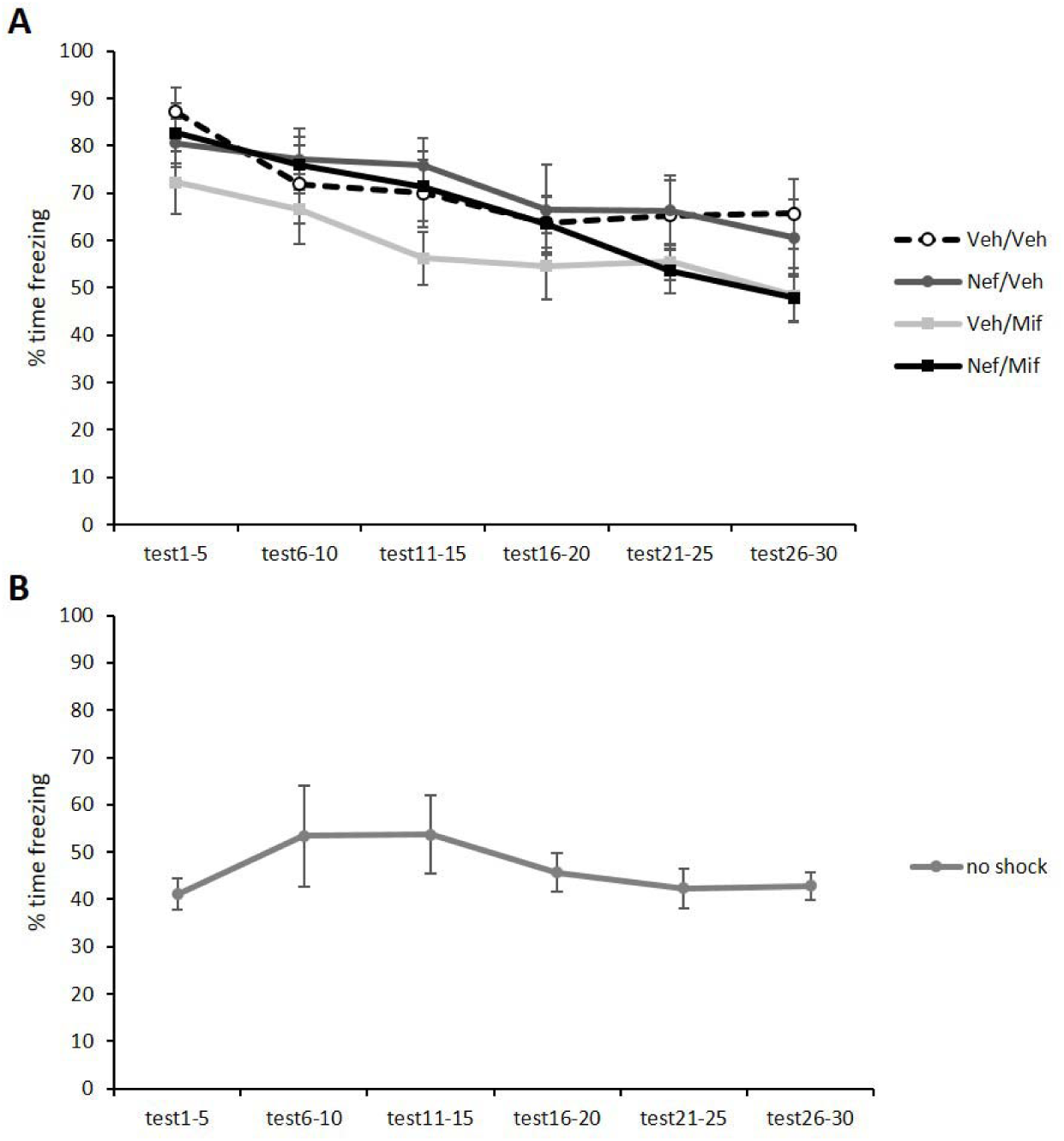
Conditioned freezing to Context B at test in groups that previously received drug administration (A) and the non-conditioned no shock control (B). Mifepristone (Mif) injected immediately after Context A memory reactivation resulted in impaired freezing to Context B at test. The evidence for the impairment was greater when reactivation was preceded by vehicle, rather than nefiracetam (Nef), injection. Data presented as mean ± SEM (n = 8 per group).

When comparing each group to the non-conditioned control, freezing was consistently higher across the session in all groups expect the Vehicle-Mifepristone group. There was a main effect of Group (F(4,35)=7.44, p<0.001, η^2^_p_=0.46, BF_inc_=107.4), with no Group x Bin interaction (F(9.7,84.8)=1.12, p=0.36, η^2^_p_=0.11, BF_inc_=0.49). Post-hoc pairwise comparisons on the main effect of Group showed that the non-conditioned group froze lower than each of the other groups (p’s<0.011, BF_10_’s>16440), apart from the Vehicle-Mifepristone group (p=0.19, but BF_10_=50.5). However, the Vehicle-Mifepristone group did not differ from the other groups (p’s>0.13, but BF_10_’s varied: 0.96 [Nef/Mif], 19.4 [Nef/Veh] & 67.3 [Veh/Veh]).

The present results show an effect of mifepristone administered immediately after reactivation of Context A fear memory to reduce generalised freezing to a different Context B. This was observed despite there being no evidence for an impairment in contextual freezing in Context A. This apparently-normal expression of contextual fear in Context A would typically be interpreted as a lack of reconsolidation impairment. However, the evidence for behavioural differences under different (Context B) test conditions suggests that mifepristone did have a subtle disruptive effect on the contextual fear memory.

The conditioning session in Context B took place 14 days after conditioning in Context A. Generalization of contextual fear is typically observed with increasing conditioning-to-test intervals of 14 days or more in rodents and is thought to reflect poorer context memory precision (Jasnow et al. 2017). The timecourse of the emergence of generalization is similar to that of systems consolidation (Squire and Alvarez 1995) or memory transformation (Winocur et al. 2007), both of which acknowledge the dependence of older contextual fear memories upon cortical regions. Interestingly, 6 hours after reactivation of a 30-day old contextual fear memory, inhibition of the anterior cingulate cortex impaired generalized fear expression but not fear expression to the training context (Einarsson et al. 2015). Moreover, there is evidence that weaker contextual fear memories display less generalization (Poulos et al. 2016). While we cannot fully explain why post-reactivation mifepristone resulted in disrupted context fear generalisation, but with intact context fear expression, this pattern is consistent with a form of memory impairment. Indeed, it is not unusual for apparently-normal behaviour to mask underlying impairments that revealed in alternative test settings. For example, PKM-ζ-null mice displayed impairments in place memory, but only under conditions of increased cognitive demand (Tsokas et al. 2016). Therefore, our results are likely to reflect a subtle manifestation of reconsolidation impairment by mifepristone, which is consistent with previous demonstrations that mifepristone impairs fear memory reconsolidation (Jin et al. 2007; Pitman et al. 2011; Flavell and Lee 2019).

The generalisation of contextual fear in the present study did not allow an assessment of stress-enhanced fear learning (Rau et al. 2005), as the learning to Context B is confounded by the differing generalised baseline freezing. Nevertheless, the persistence of the deficit through conditioning to the test in Context B provides evidence that footshock re-exposure did not reinstate the impairment in generalized freezing. Such a lack of reinstatement is typically interpreted as being consistent with an impairment in reconsolidation (Duvarci and Nader 2004).

An alternative interpretation is that mifepristone enhanced the precision of the Context A fear memory, limiting generalisation to Context B. However, memory reactivation alone has been shown to maintain the precision of contextual fear memories via memory reconsolidation (De Oliveira Alvares et al. 2013), and so such an interpretation would have to conclude that post-reactivation mifepristone enhances reconsolidation.

Returning to the reconsolidation impairment interpretation, a surprising conclusion is that the strong Context A fear memory appears to destabilize following a relatively brief context re-exposure session of 5 minutes and without the need for additional pharmacological treatment. This contrasts with previous evidence that strongly conditioned contextual fear memories in mice were not destabilized by a 5-min reactivation session following conditioning with 3 footshocks (Suzuki et al. 2004). However, the evaluation of successful destabilization was conducted in a test of freezing to the conditioned context, and so the results of Suzuki et al (2004) remain consistent with our present observations. Nevertheless, it should also be noted that we observed a significant and marked decline in freezing from the reactivation session to the test in Context A, which might be argued to be inconsistent with a strongly-learned fear memory. Therefore, it is possible that despite our multiple footshock conditioning procedure and high freezing at the reactivation session, the contextual fear memory was in fact not strong enough to produce a boundary condition on reconsolidation. Conversely, it may be that boundary conditions on reconsolidation are, in fact, more subtle than suggested by previous literature (Wideman et al. 2018) and might influence the quantitative extent or qualitative nature of memory deficits.

Here, the combination of pre-reactivation nefiracetam and post-reactivation mifepristone did not impair freezing to Context A or generalisation to Context B. While there was no strong evidence that nefiracetam actually reversed the disruptive effect of mifepristone, it remains clear that nefiracetam did not enhance the destabilization of the contextual fear memory; although it remains possible that the predicted results might have been observed under different parametric conditions. This is in clear contrast to nefiracetam’s facilitative effect on cued fear memory destabilization (Flavell and Lee 2019). One potential explanation is informed by previous observations that strong cued fear conditioning downregulates GluN2B receptor expression in the amygdala (Wang et al. 2009). As GluN2B-containing NMDA receptors are necessary for memory destabilization (Ben Mamou et al. 2006; Milton et al. 2013), this downregulation may account for the transient inhibition of memory destabilization that occurs for more than 7 days after strong fear conditioning (Wang et al. 2009).

Normalisation of the downregulation accompanies the return of memory destabilization by 30 days after conditioning. As we have previously argued (Flavell and Lee 2019), the functional mechanism of action of nefiracetam to enhance memory destabilization is likely the increase of NMDA receptor currents via interaction with the glycine binding site (Moriguchi et al. 2003). Therefore, downregulation of GluN2B receptors would be expected to limit the beneficial impact of nefiracetam on memory destabilization. This does leave the question of how contextual fear memory destabilization can proceed in spite of downregulated amygdala GluN2B receptors. Contextual fear memories, however, have an arguably expanded critical neural circuitry that includes the dorsal hippocampus (Chaaya et al. 2018) and anterior cingulate cortex (Frankland et al. 2004). As a result, the disruptive effect of mifepristone in the current study may have the dorsal hippocampus as its locus of action, compared to a likely amygdala locus of action for the effects of mifepristone on cued fear memory reconsolidation (Jin et al. 2007). Consistent with such an interpretation, mifepristone has been shown to impair the reconsolidation of hippocampal-dependent memories (Nikzad et al. 2011; Achterberg et al. 2014). Alternatively, mifepristone might have present effects in the anterior cingulate cortex, which would be consistent with the selective effect on generalized context fear expression (Einarsson et al. 2015).

In summary, post-reactivation mifepristone appears to impair the reconsolidation of strongly-conditioned contextual fear memories without the need for pharmacological enhancement of memory destabilization. Moreover, the addition of pre-reactivation nefiracetam may limit the efficacy of mifepristone. Therefore, it remains unclear whether the dual treatment approach of enhancing destabilization and impairing reconsolidation with nefiracetam and mifepristone, respectively (Flavell and Lee 2019), is of potential clinical benefit when translated to intensely-earned fear/traumatic memories. An additional implication of the current results is the vulnerability to over-interpretation of a single test of memory expression. Apparently-normal behaviour may mask underlying memory impairments, and/or may be a result of specific parameters used within the experiment. Adding multiple within-subject tests of behaviour may have value, as long as their interpretation is treated with appropriate statistical care.

## Acknowledgments

The authors would like to thank David Barber for technical support. The authors have no conflicts of interest. The experiments comply with the United Kingdom Animals (Scientific Procedures) Act 1986, Amendment Regulations 2012.

